# Higher seed yield through selection for reduced seed shattering in Italian ryegrass (*Lolium multiflorum* Lam.)

**DOI:** 10.1101/2023.12.01.569550

**Authors:** Jenny Kiesbauer, Roland Kölliker, Maria Hug, Meril Sindelar, Linda Helene Schlatter, Jonathan Ohnmacht, Bruno Studer, Christoph Grieder

## Abstract

Seed shattering, i.e., the loss of seeds at ripening stage shortly before or during seed harvest, is strongly reducing seed yield in Italian ryegrass (*Lolium multiflorum* Lam.). The aim of this study was to evaluate the possibility to reduce seed shattering within breeding germplasm via recurrent phenotypic selection on spaced plants. Starting from a founder population of 300 plants serving as F_0_ population, two cycles of phenotypic selection for high and low seed shattering were applied and compared to randomly selected individuals on spaced plant level and in plot trials. Comparison of the five resulting populations in a spaced plant trial revealed a significant effect of selection, with lowest seed shattering (15.3%) observed in the population selected twice for decreased shattering (15.3%) and highest seed shattering (47.9%) for the population selected twice for increased shattering. The same ranking of the five F_2_ populations was observed in a subsequent trial with sown plots. Thus, using the method presented here, recurrent selection on single spaced plants allows to efficiently reduce seed shattering and, therefore, to increase seed yield in swards.

## 1 INTRODUCTION

Italian ryegrass (*Lolium multiflorum* Lam.) is one of the most abundant grass species in temperate grassland used for forage production. It is an annual species, valued for its high biomass yield and fast ground cover in intensive, short term grassland management systems (Humphreys et al., 2010). Italian ryegrass is an obligate outcrossing species, resulting in a high degree of heterozygosity and a large genetic variation within natural and breeding populations. Breeding of Italian ryegrass is mainly based on recurrent phenotypic selection on spaced plants, followed by cultivar development via open pollination in a polycross and testing of cultivar candidates in plot trials (Conaghan & Casler, 2011). One limitation of this procedure is that phenotypic observations for complex traits on spaced plants cannot always be directly transferred to sward conditions as observed in a plot (Casler et al., 1996). For example, traits related to seed yield, such as number of fertile tillers or spike length, phenotyped on spaced plants only poorly correlated with the results from plot trials (Elgersma, 1990b). In general, prerequisites for successful phenotypic selection are a high heritability of the trait of interest, adequate phenotyping methods and a large phenotypic and genotypic variation within the population. The main breeding targets in Italian ryegrass are biomass yield, digestibility and palatability, disease resistance and persistence (Lüscher et al., 2019). Since low seed yield makes seed production more expensive, seed yield is also a crucial trait for a cultivar to be successful on the market (Boelt & Studer, 2010).

The potential seed yield is defined as the total number of ovules per area present at flowering time (Falcinelli, 1999). Many factors such as unsuccessful pollination and fertilization, abortion of seeds during development as well as seed shattering (i.e., the loss of seeds before harvest) can lead to realized seed yields that are substantially lower than the potential seed yield (Boelt & Studer, 2010; Marshall, 1985). Higher seed yielding varieties may be achieved by breeding for an improved potential seed yield, e.g., via increasing numbers of spikes per area, spikelets per spikes, or flowers per spikelet. Alternatively, factors leading to reduced realized seed yield as discussed above may be tackled. In Italian ryegrass, seed losses of up to 54% due to shattering have been observed (Maity et al., 2021), and even higher rates of up to 78% were observed for perennial ryegrass (*L. perenne* L.) (Tubbs & Chastain, 2022). Therefore, seed shattering seems to be a relevant breeding target to develop Italian ryegrass cultivars with improved seed yield.

Several studies showed considerable genetic variation for seed shattering among different ryegrass accessions and cultivars, indicating the potential for improving the trait through recurrent phenotypic selection (Elgersma, 1990a; Harun & Bean, 1979; Hides et al., 1993; Tubbs & Chastain, 2022). Phenotyping of seed shattering, usually defined as the proportion of seeds lost from the total amount of seeds produced (lost and non-lost), has so far mainly relied on measurements on spaced plants. In perennial ryegrass, seed shattering was measured by rolling three spikes per plant over a steel bar and calculating the percentage of seeds shattered at a defined plant maturity stage (Tubbs & Chastain, 2022). Another approach consists in bagging inflorescences or parts of it after the end of flowering, and to determine the percentage of seeds fallen off until seed harvest (Kavka et al., 2023). Further, high throughput phenotyping may be possible using an imaging pipeline for the description of spike architecture, which was found to be associated with seed shattering (Barreto Ortiz et al., 2020; Tubbs & Chastain, 2022).

Usually, grass breeding starts with the selection of single plants from large breeding populations in the spaced plant nursery. The selection could be made either directly based on the phenotypic data of the trait or by indirect selection, where the breeder improves the trait by selection for a secondary trait. A secondary trait could be beneficial if the phenotyping of the trait of interest is time consuming or not even possible. If an efficient and reliable phenotyping method is available, high selection intensities can be achieved at this stage for a like seed shattering. Further, to achieve a high selection efficiency, the genetic correlation between the secondary traits and the trait of interest as well as the heritability of the traits need to be high (Gallais, 1984).

On the other hand, seed production is usually done in swards and needs, therefore, to be tested on a plot level. Hence, a sufficient correlation of trait expression between spaced plant and sward level is a prerequisite for selection towards reduced shattering in spaced plants to be effective. Moreover, in an optimal selection system, undesired indirect selection affecting other traits like maturity should be avoided. To date, information on the efficiency of selection for reduced shattering in ryegrasses is still scarce. Employing a phenotyping system based on bagging inflorescences and standardized determination of harvest time, the objectives of this study were to (1) conduct a selection experiment to evaluate the response to recurrent phenotypic selection for this economically important trait, (2) test whether this selection on spaced plants also affects seed shattering and yield in swards, and (3) look for correlated traits that could be used as potential targets for indirect selection.

## 2 MATERIALS AND METHODS

Starting from a founder population, this study is based on two cycles of divergent selection towards reduced and increased seed shattering on spaced plants, respectively. This resulted in five F_2_ populations differently selected for seed shattering: F_2_ neutral (no selection for seed shattering), F_2_+ (one cycle of positive selection [low seed shattering]), F_2_++ (two cycles of positive selection), F_2_- (one cycle of negative selection [high seed shattering]) and F_2_-- (two cycles of negative selection; Figure 1). In comparative trials on a spaced plant and sward level, these five populations were phenotyped for different traits to assess the effect of selection.

**FIGURE 1.**
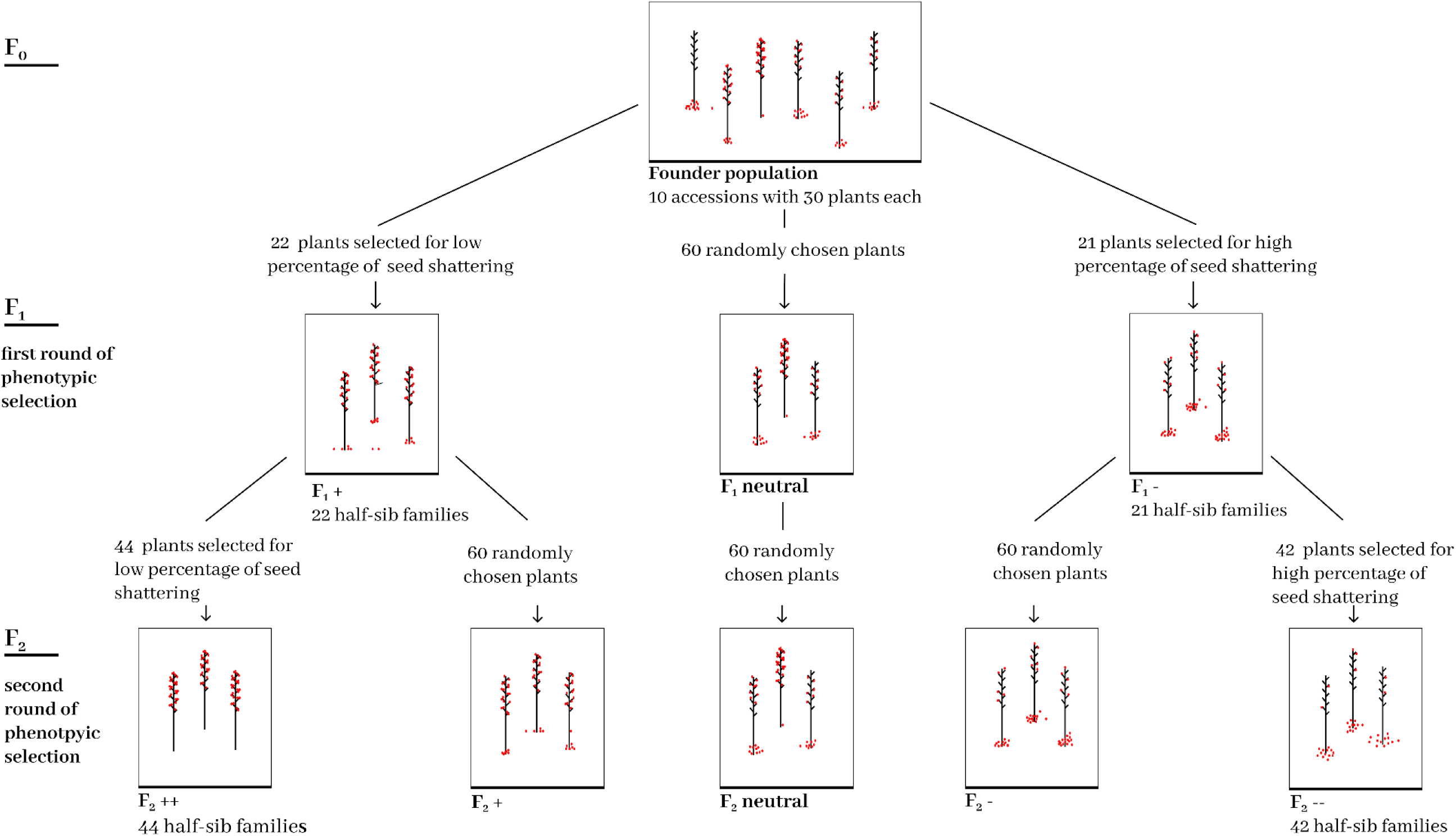
Scheme of the selection experiment. Starting from a founder population (F_0_), a first phenotypic selection was made based on spaced plants in the field and plants with contrasting levels of seed shattering were open pollinated to produce three F_1_ populations (F_1_+, F_1_-, and F_1_ neutral). F_1_ plants were again selected for *low and high* seed shattering, giving rise to two F_2_ populations selected twice for low (F_2_++) and high seed shattering (F_2_--). respectively. In addition, plants were randomly chosen from the two selected F_1_ populations resulting in two additional F_2_ populations with only one cycle of positive or negative selection (F_2_+ and F_2_-). An unselected F_2_ population (F_2_ neutral) was obtained by open pollination of randomly chosen plants over two generations.

### 2.1 FIRST CYCLE OF SELECTION

The founder population serving as starting point for selection (F_0_), consisted of ten synthetic populations (SYN2) from the Agroscope breeding program of diploid Italian ryegrass (Table 1). Each SYN2 originated from a polycross of variable size where the seeds harvested from the polycross (SYN1) were multiplied for another generation in the field and harvested in bulk. Two of the ten SYN2 are currently registered as cultivars (’Xanthia‘ and ’Bipes’).

**TABLE 1.** Synthetic populations (SYN2) contributing to the F_0_ generation, including cultivar name for those officially registered, number of genotypes contributing to the polycross (PC size) and number of F_0_ plants selected as parents for the positive (low seed shattering, F_0_+) and negative (high seed shattering, F_0_-) selection in the first cycle.

| Population | Cultivar name | PC size | Number of plants |  |
| --- | --- | --- | --- | --- |
| | | | $F_{0+}$ | $F_{0-}$ |
| LI0615 | Xanthia | 23 | 2 | 2 |
| LI0835 | n.a. | 18 | 3 | 2 |
| LI1015 | n.a. | 11 | 3 | 2 |
| LI1105 | n.a. | 21 | 2 | 2 |
| LI1115 | n.a. | 13 | 1 | 3 |
| LI1135 | Bipes | 8 | 3 | 2 |
| LI1235 | n.a. | 14 | 2 | 2 |
| LI1245 | n.a. | 10 | 2 | 3 |
| LI1255 | n.a. | 11 | 3 | 2 |
| LI1265 | n.a. | 12 | 2 | 3 |

For each of the ten SYN2, 30 genotypes, each one represented by one plant, were transplanted to the field in August 2016 at the Agroscope research station in Zurich-Reckenholz, Switzerland. A typical breeding nursery design was used, with ten plants of the same SYN2 being planted in a row (hereafter denoted as nursery-row). The planting distance was 40 cm within and 50 cm between nursery-rows. Three blocks of 10 nursery-rows (one per SYN2) were formed. The nursery trial was managed according to standard practices. In 2017, plants were cut once at the beginning of May. In the second growth, plant vigor was rated on a scale from 1 (very good) to 9 (very poor) and seed shattering was assessed according to the standard protocol described in section 2.3.1, with the only exception that seeds were harvested on two separate dates (for early and late flowering plants). Twenty-three genotypes with low-seed shattering and good vigor (Table 1; F_0_+) and 23 genotypes with high seed shattering and good vigor (F_0_-) were selected as parents of the F_1_ populations. For both selections, culms cut shortly before flowering during the first growth in 2018 were allowed to pollinate in two separate isolations in the greenhouse for seed production. F_1_ seeds were harvested separately for each maternal F_0_ plant, whereby one genotype of the positive selection and two genotypes of the negative selection did not set seed, resulting in 22 and 21 half-sib (hs) families forming the F_1_+ and F_1_-population, respectively (Figure 1).

To establish the F_1_ neutral population, 60 randomly chosen plants of the F_0_ population were cut from the field and open pollinated in the greenhouse.

### 2.2 SECOND CYCLE OF SELECTION

For all F_1_ hs-families of the F_1_+ (22) and the F_1_- (21) population, 20 genotypes, each represented by two plants, were raised in the greenhouse for three months and transplanted to the field in August 2019. The same breeding nursery design as for the first cycle of selection was used with genotypes of the same hs-family planted in two adjacent nursery-rows. An alternating pattern with two rows of F_1_+ plants followed by two rows of F_1_- plants was used. For the second replicate (i.e., the second plant per genotype) the same layout and randomization was used. In 2020, plants were cut at the beginning of May and plant vigor as well as seed shattering were again determined on the second growth. In addition, occurrence of stem rust (caused by *Puccinia graminis* subsp. *graminicola*) was rated on a scale from 1 (no rust occurrence) to 9 (very heavy rust infestation) at time of seed harvest according to Schubiger and Boller (2016).

Two genotypes of each F_1_+ hs-families (44 in total) with low seed shattering, acceptable vigor (< 3) and no or low stem rust occurrence (F_1_++), and two genotypes of each F_1_- hs- family (42 in total), with high seed shattering, acceptable vigor and no or low stem rust occurrence (F_1_--), were selected as parents for the respective F_2_ populations. One plant per selected genotype was dug out from the field in September 2020. Potted plants were kept outside for vernalization and were transferred to the greenhouse in January 2021. These plants were allowed to flower (separate chamber per selection) and set F_2_ seeds. To generate seeds of the F_2_+ and F_2_- populations, approximately 60 randomly chosen culms of F_1_+ and F_1_- plants, respectively, were cut from the field and open pollinated in the greenhouse. To establish the F_2_ neutral population, approximately 60 randomly chosen plants of the F_1_ neutral population were cut from the field and open pollinated in the greenhouse.

### 2.3 COMPARATIVE TRIALS

During the field season 2021 and 2022, the populations divergently selected for seed shattering (F_2_++, F_2_+, F_2_ neutral, F_2_-, F_2_--; Table 2) were compared in two field experiments with spaced plants and sown rows.

**TABLE 2.** Origin of the five F_2_ populations (F_2_ neutral, F_2_+, F_2_++, F_2_-, F_2_--). The type indicates whether half sib families (hs-fam.) of the F_1_ population or randomly chosen (bulk) plants of the F_1_ population were used to generate the F_2_ population.

| Population | Gen. | Type | Origin |
| --- | --- | --- | --- |
| $F_2--$ | $F_2$ | 42 $F_1$ hs-fam. | Seeds harvested on 42 $F_1-$ plants selected for high seed shattering in the second cycle (open pollination in the greenhouse among potted plants) |
| $F_2-$ | $F_2$ | bulk | Seeds harvested on ~ 60 randomly chosen $F_1-$ plants (open pollination in the greenhouse among culms cut from the field) |
| $F_2$ neutral | $F_2$ | bulk | Seeds harvested on ~ 60 randomly chosen $F_1$ neutral plants, (open pollination in the greenhouse among culms cut from the field) |
| $F_2+$ | $F_2$ | bulk | Seeds harvested on ~ 60 randomly chosen $F_1+$ plants (open pollination in the greenhouse among culms cut from the field) |
| $F_2++$ | $F_2$ | 44 $F_1$ hs-fam. | Seeds harvested on 44 $F_1+$ plants, selected for low seed shattering in second cycle (open pollination in the greenhouse among potted plants) |

#### 2.3.1 SPACED PLANT TRIAL

To compare spaced plants with plot data, a field trial was installed in August 2021 at the Agroscope research station in Zurich-Reckenholz, Switzerland. For each of the five F_2_ populations, 80 genotypes, each represented by one plant previously raised in the greenhouse for two months, were transplanted to the field. The same planting scheme as for the first selection cycle was used. Nursery-rows containing different populations were randomly distributed in the field.

Phenotyping was performed on single spaced plants. In 2022, heading date (determined in days after first of April) and plant vigor of the first growth was assessed. After completion of heading, plants were cut. For the second growth, plant vigor and heading date were again determined. Additionally, begin of flowering (also determined in days after first of April) was assessed. The occurrence of stem rust was rated on a scale from 1 to 9 (see above). The occurrence of late culms, i.e., culms that appear later and grow much shorter, was also assessed on a scale from 1 (no late culms) to 9 (numerous late culms). To determine seed shattering, the complete inflorescence of each plant was bagged after termination of flowering into a perforated plastic bag (Sealed Air, Cryovac, Charlotte, USA 330mm x 750mm) and tied to a bamboo stick. Time of harvest was determined using the temperature sum (based on the average daily temperature sum 5 cm above ground from a nearby weather station). The day of harvest of the earliest flowering plant was determined manually according to experience, all other plants were harvested on the day when the temperature sum accumulated since their start of flowering reached the same value as for the earliest flowering plant. At harvesting, the bagged inflorescences were removed from the plant and gently shaken three times by hand to ensure that seeds hanging loose but not yet shattered fall off the spikes into the bag. The bag was then opened, the inflorescence removed and transferred to a second perforated plastic bag. Shattered seeds (still in the first bag) and non-shattered seeds (in the second bag together with the inflorescence) were then dried in a drying cabinet under constant air flow at room temperature for three days. Non-shattered seeds were manually removed from the inflorescences. During cleaning of shattered and non-shattered seeds, empty seeds and debries were removed from fully developed seeds using an airflow-separator (Saugluft-Stufensichter T2, Baumann Saatzuchbedarf, Waldenburg, Germany). Weight of cleaned shattered and cleaned non-shattered seed per plant was determined using a digital scale (New Classic MS; Mettler Toledo; Columbus Ohio, accuracy 0.01g). Seed shattering (%) was then calculated following the formula.

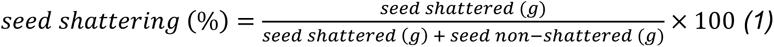

#### 2.3.2 PLOT TRIAL

Plot trials with the five F_2_ populations were sown in August 2021 at the three locations in Zürich-Reckenholz (47.4301°N, 8.5235°E), Rümlang (47.4380°N, 8.5290°E) and Oensingen (47.2840°N, 7.7321°E). Rows of 2.5 m length were sown by hand with a distance of 0.5 m between rows. Two rows next to each other formed a plot, and four plots were sown per F_2_ population at each location, arranged in a randomized complete block design. This resulted in 20 double-row plots per location and 60 double-row plots in total. To avoid border effects, additional rows were sown at the edge of each trial. Trials were managed according to standard procedures.

In 2022, rows were cut in early May before heading and plastic foil strips of 0.45 m x 2.0 m were placed between the two rows at the center of a double-row plot shortly thereafter. Rows were then allowed to grow, flower and set seeds. At each location, plots were harvested at two different dates, i.e., early vs. late harvesting. Harvest dates were on 30 June and on 6 July for locations Zürich-Reckenholz, and Rümlang and on 4 July and on 8 July for location Oensingen. At each harvest date, culms with inflorescences of complete double-row plots were cut, put into fabric bags and put to the drying cabinet at constant airflow for 3 days. In addition, the seeds shattered onto the plastic foil strips within double-row plots were collected. Further determination of seed shattering, i.e., removal of non-shattered seeds from inflorescences, seed cleaning and calculation of proportion of shattered seeds, was done as described for the spaced plant trial. In addition, plant vigor was rated on a one to nine scale as described for spaced plants. Heading date during the second growth was also assessed, but as variance within plots was larger than variation among plots, no differences could be determined (all plots heading at the same day).

### 2.4 DATA ANALYSIS

All statistical analysis were conducted with R v4.1.2 within RStudio v4.0.5 (R Core Team, 2020; RStudio Team, 2021) using standard functions for different calculations, functions “lm” and “anova” for classical ANOVA and function “lmer” from the package “lme4” for mixed model analyses (Bates et al., 2015).

#### 2.4.1 Spaced plants

The spaced plant trial followed a split-plot design, where nursery-rows with ten plants of the same selection formed the main-plot stratum and single plants formed the sub-plot stratum. To account for this non completely randomized design, the following mixed model was used:

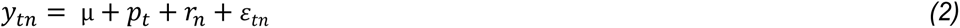

where *y_tn_* represents the observation for trait *y* on a single plant basis, *µ* denotes the overall mean, *p_t_* is the effect of population t, *r_n_* the effect of nursery-row n, and *ε_tn_* the residual error. Factor *p_t_* was taken as fixed, whereas *r_n_* was random. Mean values per population were calculated and F_2_++, F_2_+, F_2_--, F_2_- were each compared to F_2_ neutral using Dunnett’s test for multiple comparison. Realized heritability (*h^2^*) was calculated based on the breeder’s equation as:

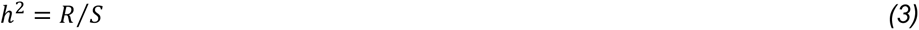

where R is the realized response to selection and S the selection differential. Realized heritability could only be calculated for the second selection cycle, i.e., from F_1_+ to F_2_++ and from F_1_- to F_2_--. S was calculated based on data assessed from the trial with F_1_ plants. For the second positive selection step, S was defined as the difference between the mean performance of all F_1_+ plants to the mean performance of the F_1_+ selected as parents for the F_2_++ population. For the second negative selection step, S was defined as the difference between the mean performance of all F_1_- plants to the mean performance of the F_1_- plants selected as parents for the F_2_-- population. R was calculated based on data assessed from the comparative spaced plant trial with F_2_ plants in 2022. For the second positive selection step, R was calculated as mean of all F_2_++ plants to the mean of all F_2_+ plants. For the second negative selection step, R was calculated as mean of all F_2_-- plants to the mean of all F_2_- plants.

#### 2.4.2 Sown plots

Analysis of variance (ANOVA) for the traits assessed in the sown row trial was performed using general linear models. In a pre-analysis, all factors, i.e., population, location and harvesting as well as all of their possible interactions were tested in a fully factorial model. As the triple interaction as well as the population-by-location interaction did not show a significant effect for any of the analyzed traits (data not shown), the following model was used:

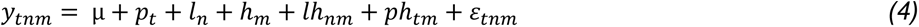

where *y_tnm_* represents the observation for trait *y* on a (double-row) plot basis, *µ* denotes the overall mean, *p_t_* is the effect of population t, *l_n_* the effect of location n, *h_m_* the effect of harvesting timepoint m, *lh_nm_* the interaction between location n and harvesting timepoint m, *ph_tm_* the interaction between population t and harvesting timepoint m, and *ε_tnm_* the residual error. Correlations between the means of the five space plant populations and the means of the early and the late plot trials were calculated as Spearman’s rank correlations.

## 3 RESULTS

Seed shattering as observed in the spaced plant trial showed a clear effect of selection. The highest mean percentage of seed shattering was observed in the F_2_-- (47.2%), followed by the F_2_- (41.9%), F_2_ neutral (38.7%), F_2_+ (22.5%) and F_2_++ (15.3%) population (Figure 2). Analysis of variance indicated a significant effect of the population on seed shattering (*p* < 0.001, data not shown). A comparison of the F_2_+, F_2_++, F_2_-, F_2_-- to the F_2_ neutral population with Dunnett’s test revealed that already one cycle of selection significantly decreased seed shattering (F_2_ neutral vs. F_2_+, *p* < 0.001) and this difference even increased with the second selection cycle (F_2_ neutral vs. F_2_++, Figure 2). On the other hand, the selection for higher seed shattering did not increase seed shattering after one cycle of selection (F_2_ neutral vs. F_2_-, *p* > 0.05), and the difference was still not significant after a second cycle of selection (F_2_ neutral vs. F_2_--, *p* > 0.05).

**FIGURE 2.**
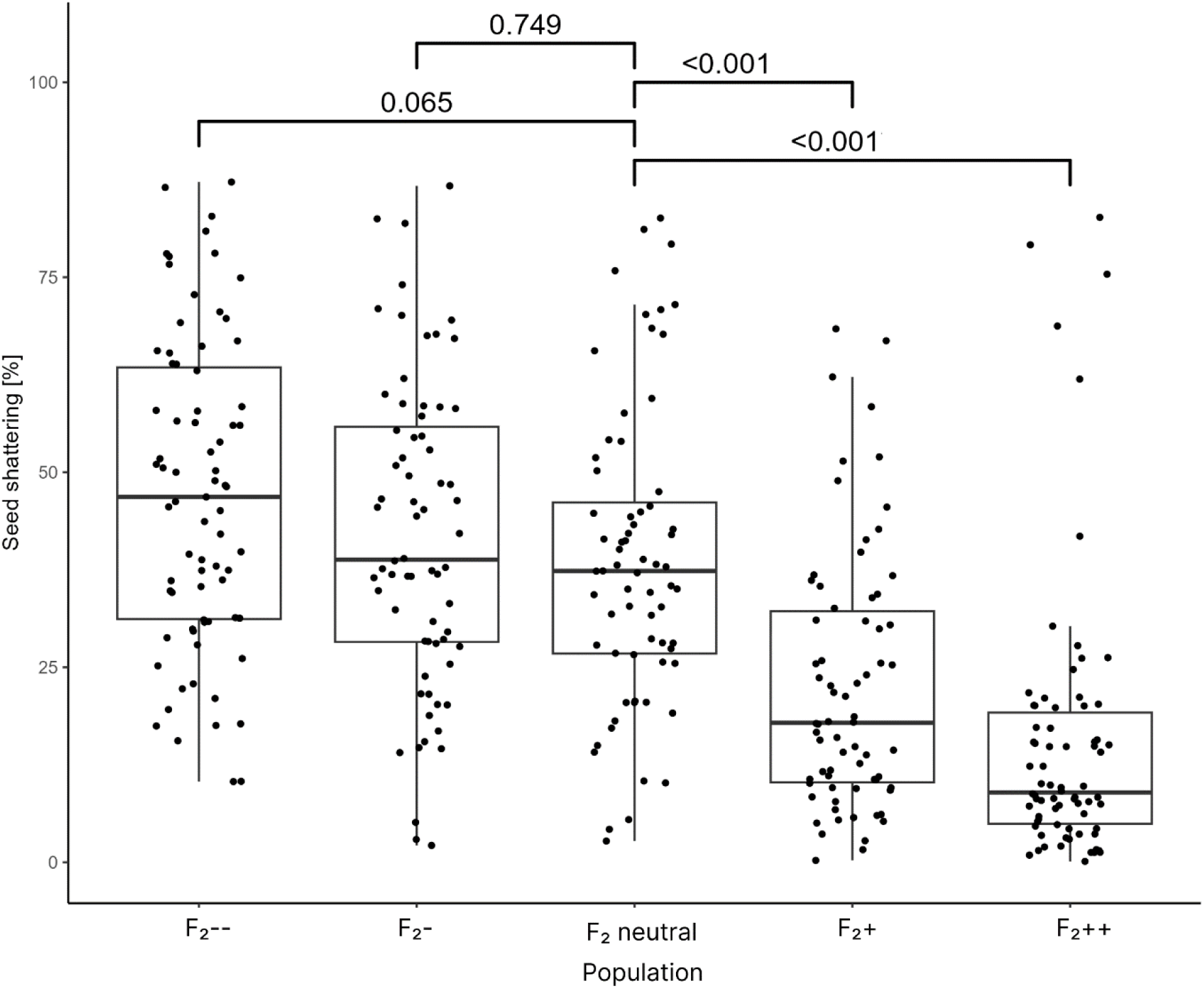
Seed shattering as assessed on spaced plants of five Italian ryegrass populations: F_2_-- was selected twice for high seed shattering (double negative selection), F_2_- was selected once for high seed shattering (negative selection), F_2_ neutral was not selected, F_2_+ was selected once for low seed shattering (positive selection) and F_2_++ was selected twice for low seed shattering (double positive selection). Bars with corresponding numbers give the *p* value of the comparison with the F_2_ neutral population.

Calculation of realized heritability indicated low to moderate values of h^2^ = 0.20 and 0.32 for the second negative and second positive selection cycle, respectively (Table 3). Although the selection differentials (S) were of nearly equal size (24.4 for positive and 23.0 for negative selection), the lower heritability for the negative selection resulted in a pronouncedly reduced response to selection (R) compared to the positive selection.

**TABLE 3.** Realized heritability (*h^2^*) for the selection steps leading to the F_2_ populations F_2_++ and F_2_--. The selection differential (*S*) is calculated as the difference between the mean of all and the mean of the selected genotypes from the corresponding F_1_ populations (F_1_+ and F_1_-). The response to selection (*R*) is calculated as the difference between the means of the F_2_++ and F_2_+ (or F_2_-- and F_2_-) population from the comparative spaced plant trial.

| Selection | Seed shattering [%]<br>in $F_1$ generation | | | Seed shattering [%]<br>in $F_2$ generation | | | $h^2$ |
| --- | --- | --- | --- | --- | --- | --- | --- |
| | All | Selected | $S$ | Single | Double | $R$ | |
| $F_{1+}$ to $F_{2++}$ | 32.3 | 7.9 | -24.4 | 22.6 | 14.8 | -7.8 | 0.32 |
| $F_{1-}$ to $F_{2--}$ | 53.1 | 76.1 | 23.0 | 41.9 | 46.6 | 4.7 | 0.20 |

Seed shattering showed strongest correlations with the two components it was calculated from, i.e., the weight of shattered seeds (*r* = 0.85***) and the weight of non-shattered seeds (*r* = -0.65***, Table 4). Seed shattering only showed weak (*r* = -0.16** to -0.21**) correlations with earliness of plants as indicated by heading date of the first and second growth as well as begin of flowering (BOF). The negative correlations indicate that later maturing plants tended to show lower seed shattering. Other traits like vigor of plants, the occurrence of late culms or stem rust did have no or only a very small effect on seed shattering. Weight of shattered and non-shattered seeds showed a weak negative correlation (*r* = -0.23**), indicating that plants with a higher number of shattered seeds tended to have less non-shattered seeds.

**TABLE 4.**
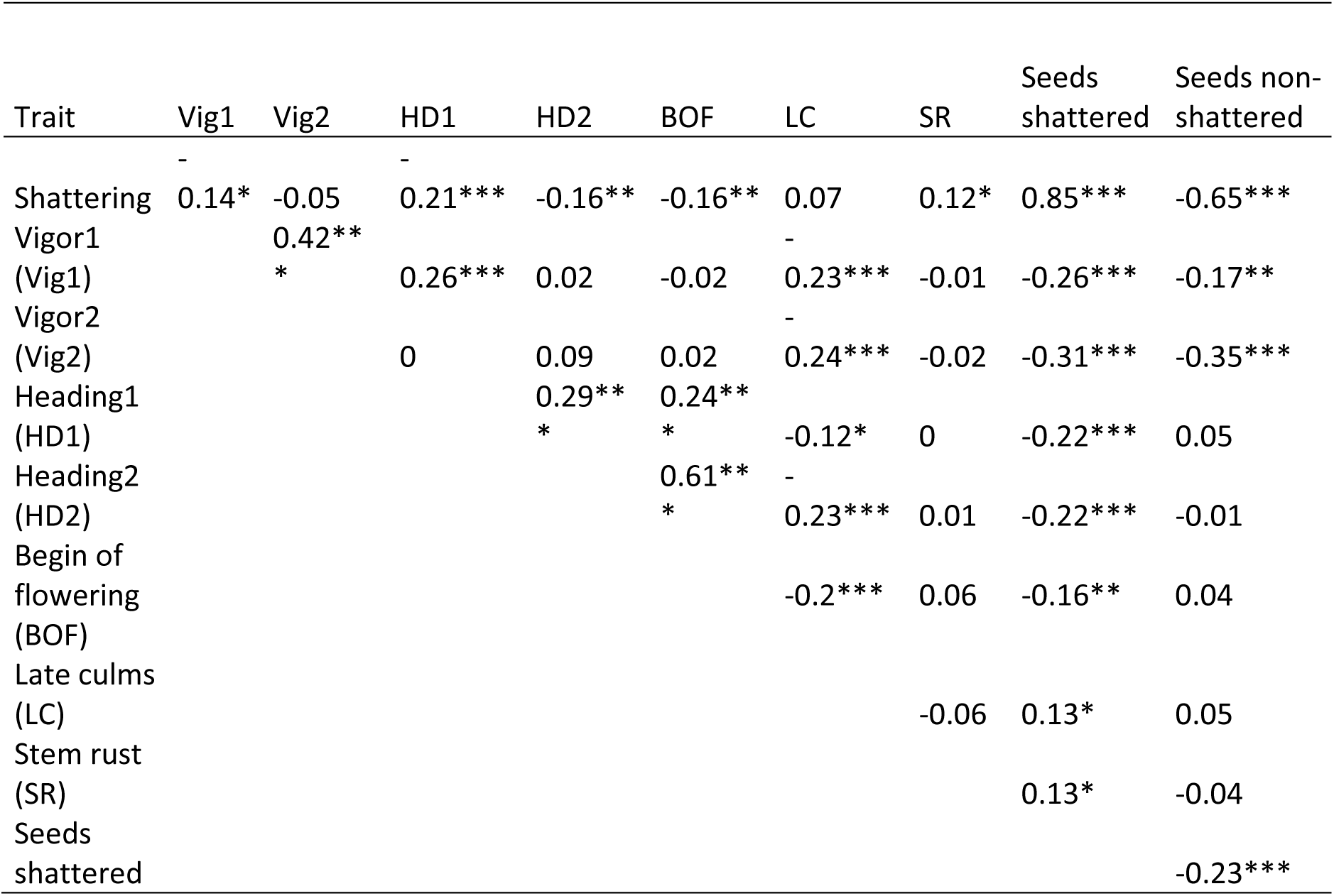
Spearman’s rank correlations among traits assessed in the comparative spaced plant trial. Vig1 = vigor observed on first growth, Vig2 = vigor observed on second growth, HD1 = heading date observed on first growth, HD2 = heading date observed on second growth, BOF = begin of flowering observed on second growth. Late culms (LC) as well as stem rust (SR) were phenotyped at harvesting date, seed shattering (%) was determined as the ratio of seeds shattered (g) over total seed weight per plant. Seeds non-shattered is the weight (g) of the seeds still on the tillers at harvesting date. Significance levels are: * = *p* value < 0.05, ** = *p* value < 0.01 and *** = *p* value < 0.001.

While seed shattering was not correlated to plant vigor, negative correlations of the weight of shattered as well as non-shattered seeds with plant vigor in the first and second growth indicated that more vigorous plants have in total more seeds, which are either being shattered or remaining on the plant. The only low to moderate correlation between heading date of the first and the second growth is largely driven by the reduced variation for this trait in the second growth and indicates that earliness from the first growth does not directly translate to the second growth. The moderate to strong correlation (*r* = 0.61***) of heading date in the second growth with BOF in the second growth indicates earlier flowering of early heading plants.

Seed shattering in sown plots was significantly affected by the type of selection, harvest time, location, as well as location-by-harvest time and population-by-harvest time interaction (Table 5). The same was the case for weight of non-shattered seeds, whereas the other two components of seed shattering, i.e., weight of shattered seeds and total seed weight, were not significantly influenced by the population-by-harvest time interaction. Plant vigor was significantly influenced by location and by harvest time, but not by the other factors or their interactions.

**TABLE 5.** Mean squares from analysis of variance (ANOVA) for different traits assessed in sown rows at three locations. Significance levels are shown as “*” = *p* value < 0.05, “**” = *p* value < 0.01 and “***” = *p* value < 0.001.

| Source of variation | df | Seed shattering (%) | Seed total (g) | Seed shattered (g) | Seed non-shattered (g) | Vigor |
| --- | --- | --- | --- | --- | --- | --- |
| Population | 4 | 254*** | 15316*** | 558*** | 21633*** | 0.06 |
| Location | 2 | 36* | 26245*** | 916*** | 17354*** | 9.8375*** |
| Harvesting | 1 | 2802*** | 115397*** | 7988*** | 184107*** | 1.35* |
| Location:Harvesting | 2 | 89*** | 3785** | 429*** | 6237*** | 0.24 |
| Population:Harvesting | 4 | 71*** | 1648 | 44 | 1683* | 0.24 |
| Residuals | 46 | 7 | 670 | 52 | 628 | 0.20 |

Seed shattering was lower for early compared to late harvesting for all five populations but ranking of the populations was the same for both harvest times (Figure 3). The highest percentage of seed shattering was observed in the F_2_-- (7.92% early, 28.44% late), followed by the F_2_- (7.34% early, 21.56% late), F_2_ neutral (6.12% early, 20.22% late), F_2_+ (4.13% early, 16.73% late) and F_2_++ (2.41% early, 9.31% late) populations (Figure 3). The difference in seed shattering between early and late harvesting continuously decreased from 20.52% in the F_2_-- population to 6.9% in the F_2_++ population, indicating that the absolute effect of the population was larger for late compared to early harvesting. Comparing the different F_2_ populations to the F_2_ neutral selection using Dunnett’s test, only the F_2_++ was significantly different (*p* = 0.010, data not shown). The correlations between seed shattering as measured in plots and seed shattering as measured in spaced plants were calculated based on mean values of the five populations. Both harvesting times showed the same correlation to the spaced plant trial (r= 1 and *p* = 0.017).

**FIGURE 3.**
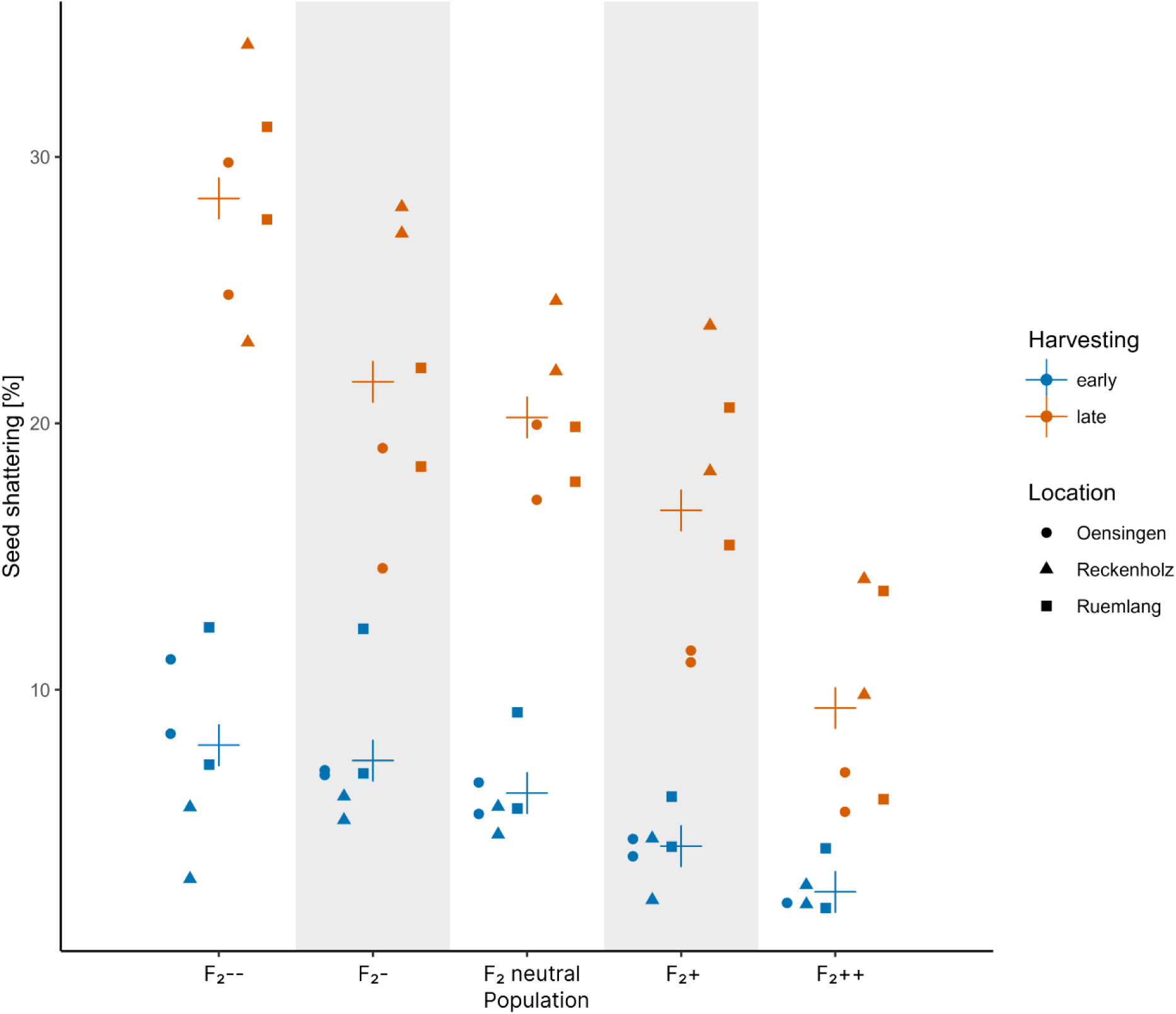
Seed shattering as determined in plots with sown rows. Blue and orange colors indicate early and late harvesting, respectively. Crosses give the average over locations per selection and harvesting time.

The harvestable seed yield in plot trials, i.e., weight of non-shattered seeds, was distinctively higher for early compared to late harvesting, but the ranking of the differently selected populations was comparable between the two harvesting timepoints (Figure 4). For late harvesting, harvestable seed yield doubled from the F_2_-- (117.5 g) to the F_2_++ population (254.9 g), with intermediate values of 149.3 g, 171.6 g and 194 g for the F_2_-, F_2_ neutral and F_2_+ selection, respectively. In comparison, differences among the populations were not as pronounced for early harvesting, where F_2_--, F_2_-, F_2_ neutral and F_2_+ populations all showed similar values for weight of non-shattered seeds (266.6 g, 264.6 g, 277.7 g and 283.4 g, respectively). Only the F_2_++ population showed a distinctively increased weight of non-shattered seeds at early harvest (349.0 g). For early harvesting, the weight of non-shattered seeds was substantially lower for location Oensingen compared to the other sites (see blue dots in Figure 4).

**FIGURE 4.**
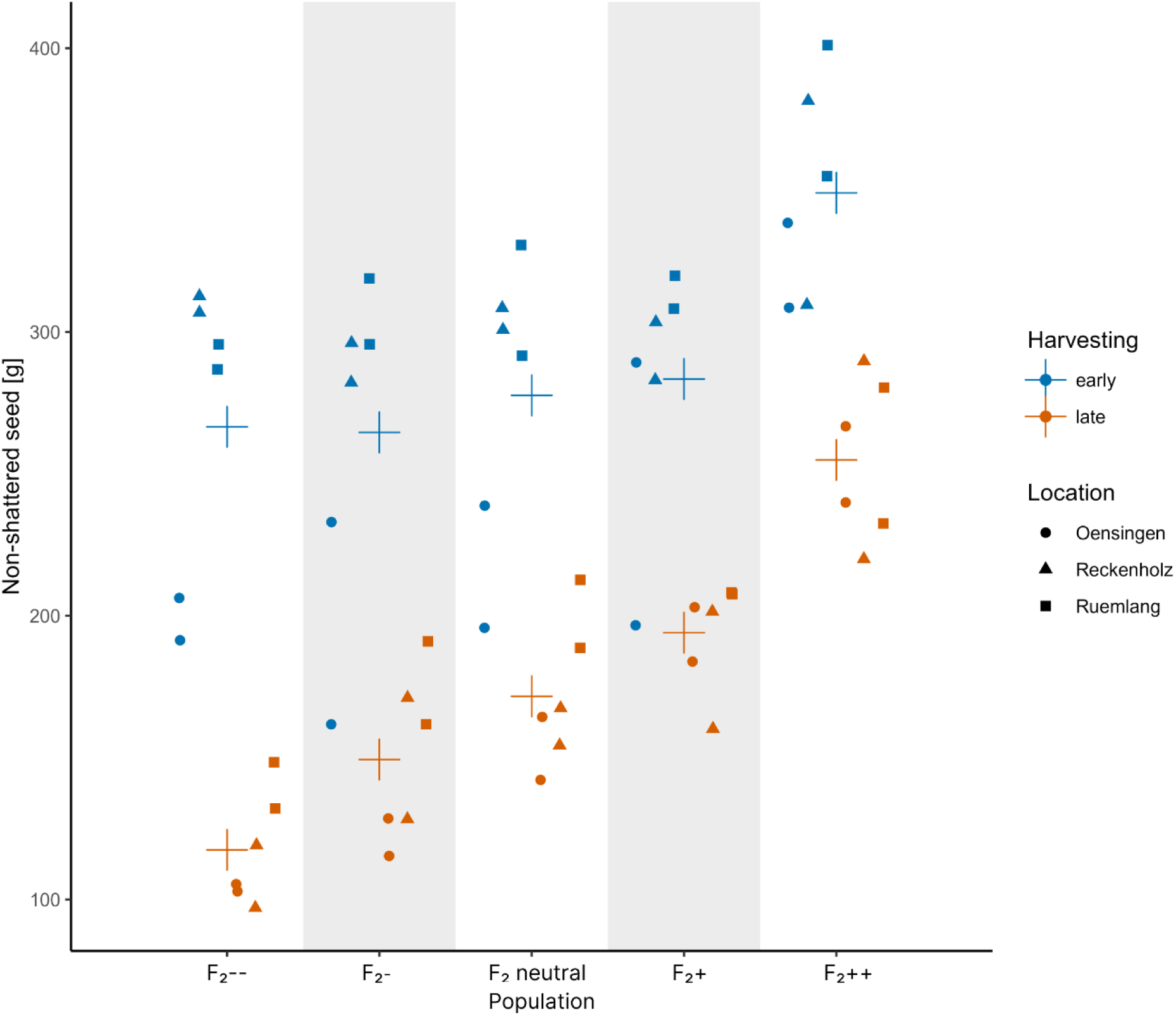
Harvestable (non-shattered) seeds as realized in plots with sown rows. Blue and orange colors indicate early and late harvesting, respectively. Crosses give the average over locations per selection type and harvesting time.

## 4 DISCUSSION

In this study, we could show that recurrent phenotypic selection for reduced seed shattering in spaced plants is very effective for Italian ryegrass. The phenotyping of spaced plants is relatively fast and easy, only a small amount of seed is required and in a second step, selection within and among families is possible (Vogel & Pedersen, 1993). The improvement of 23.4% after two cycles of recurrent selection is comparable to recurrent phenotypic selection for improving seed yield in perennial ryegrass (Marshall & Wilkins, 2003). In the first selection cycle, the selection was based on data from only one environment and one replicate per genotype grown as spaced plants. Already these phenotypic data were sufficient to realize a decrease of 16.2% in seed shattering when comparing the progenies of the F_2_ neutral and F_2_+ in the spaced plant comparative trial. The improvement was lower in the second cycle of selection (7.2%. from F_2_+ to F_2_++), although the phenotypic data was of better quality in the second selection step (two replicates per genotype in the second compared to one replicate in the first selection step). This might be partly explained by a smaller selected fraction in the first selection step (23 out of 300 = 7.7%) compared to the second selection step (42 out of 420 = 10%).

The F_0_ generation in our experiment consisted of breeding material that traces back to collections from semi-naturally occurring populations (ecotypes) (Peter-Schmid et al., 2008). This material has so far not undergone targeted selection to reduce seed shattering (data not shown). As high seed shattering in wild populations is rather an advantage to increase dispersal of seeds and, therefore, increase the spread of offspring (Dong & Wang, 2015), the shattering trait is likely to be very abundant in these materials (Piccirilli & Falcinelli, 1989). In general, selection for a trait not selected before will lead to rapid improvements in the first selection cycles, whereas after some cycles, an improvement will be harder to achieve. This would be particularly pronounced if the trait is influenced by dominant genes and the frequency of the advantageous, dominant alleles increases from low initial levels. For example in red clover, improvement of resistance to the fungal disease southern anthracnose, caused by *Colletotrichum trifolii*, revealed the highest improvement of up to 52% after one cycle of phenotypic selection in susceptible to moderately resistant cultivars (Schubiger & Grieder, 2019). After a second cycle of selection, improvement for resistance to southern anthracnose was already distinctly lower (Jacob et al., 2013). This might also explain why negative selection did not have a significant effect even after two cycles, as the frequency of the alleles leading to high shattering, already at a very high level in the F_0_ population, could not be increased any more. The realized heritability (h^2^) for the negative selection was lower than for the positive selection (Table 3), indicating that decreasing seed shattering is more effective than increasing seed shattering within these populations. Dry matter yield, one of the most important and most selected traits within forage grasses, nowadays displays an improvement of 0.17% to 0.80% per breeding cycle, depending on the species (Grieder et al., 2019). This is substantially lower compared to the rates of improvement for seed shattering we observed here. Thus, recurrent phenotypic selection as performed in our experiment with bagging inflorescences after flowering and harvesting according to sum of temperature is a valid method to decrease seed shattering in spaced plants, which is especially effective when starting from populations with high levels of shattering.

Indirect selection, where the breeder improves a primary character by selecting for a secondary character, is often used in breeding for traits that are difficult to phenotype (Gallais, 1984). Here, the employed system for phenotyping seed shattering is labor intensive and time consuming. Therefore, another, simpler trait to indirectly select for low seed shattering would be preferable. Several agronomically important traits were assessed and correlated with seed shattering. A moderate negative correlation (r = -0.65) was found between seed shattering and the weight of non-shattered seeds, indicating that selection for plants with a high number of seeds remaining on the plant could indirectly reduce seed shattering. Whether this is only the case in this population or a general observation for Italian ryegrass remains to be clarified. Selection for high seed yield (i.e., non-shattered seeds) would be an easier phenotyping method to reduce seed shattering as bagging of inflorescences is not necessary.

Relative efficiency of indirect selection, i.e., improvement of seed shattering when selecting for non-shattered seeds, can be calculated according to (Falconer & Mackay, 1996) as

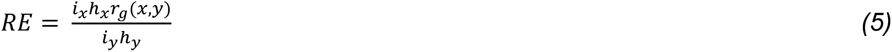

where *i* denotes the selection intensity and *h* the square root of heritability for the directly selected trait *x* (non-shattered seeds) and an indirectly selected trait *y* (seed shattering), and *r*_*g*_(*x*, *y*) is the genetic correlation between the two traits. Thus, in addition to a high *r*_*g*_(*x*, *y*), relative efficiency of indirect selection depends on the ratios *i_x_*:*i_y_* and *h_x_*:*h_y_*. The realized heritability for non-shattered seeds cannot be calculated from our data set as the selection was only done on seed shattering. Hence, no detailed statement is possible for the ratio *h_x_*:*h_y_*. However, when assuming similar heritabilities for the two traits (i.e. *h_x_*:*h_y_* ≈ 1) and the phenotypic correlation coefficient of 0.65 as a lower estimate for *r*_*g*_(*x*, *y*), indirect selection would be at least as efficient if *i_x_*:*i_y_* would be >1.54. Hence, if the faster phenotyping method for non-shattered seeds would allow to phenotype more plants and, therefore, allow for a selection intensity that is at least 50% higher, indirect selection would be effective.

Selection for a particular trait may lead to unintentional selection for another agronomically important trait. For example, a tradeoff between vegetative and reproductive growth is reported for forage grasses (Humphreys et al., 2010). On one side, Italian ryegrass as a forage crop needs to have a high vegetative biomass yield, on the other side, seed yield as a reproductive trait is also important for the successful seed multiplication of cultivars (Sampoux et al., 2011). Therefore, other agronomically important traits such as vigor, ripening time, stem rust resistance and the number of late culms were assessed for unintentional correlations with seed shattering. Low correlation coefficients observed in our study indicate that vigor and seed shattering can be selected independently from each other. Furthermore, late ripening plants tend to have less shattered seeds, which could favor a shift towards later maturity when breeding for reduced seed shattering. One possible reason could be that seeds of late maturing plants have less time to ripen and, therefore, to shatter seeds. To prevent indirect selection of late maturing genotypes, the harvesting time was determined according to the temperature sum after start of flowering. Consequently, no systematically preference for early or late breeding materials was observed within the recurrent spaced plant selection. Also in the plot trials, all the plots at a given location, independent on the selection group, headed on the same time (data not shown). Therefore, also in plot trials, no shift towards early or late maturing plants could be observed. The number of late culms did not show any effect on seed shattering. Stem rust usually develops late in the season on the panicle and florets where it may directly interfere with seed set, resulting in less viable seeds (Barker et al., 2003). Consequently, several studies reported a negative correlation between the occurrence of stem rust and seed yield (Leonard & Szabo, 2005; Pfender, 2001). Whether seed shattering is negatively affected by stem rust remains yet unclear. However, within our field trials, stem rust occurred in each environment and year. In order not to influence the selection by stem rust, only plants with no or very weak rust symptoms (score ≤ 2) were selected from all populations. Based on the correlations in spaced plants (Table 4), occurrence of stem rust only weakly increased seed shattering, mainly via increasing the weight of shattered seeds. This weak effect of stem rust occurrence on seed yield traits might be explained by a limited variation and a preponderance of other factors affecting seed yield traits. To conclude, breeding for a low seed shattering cultivar should be possible without compromising vigor, maturity or other traits. However, to obtain high seed yielding plants, breeders must select more consistently for stem rust resistant plants.

Even though seed shattering is genetically controlled, seed shattering is also strongly influenced by agronomical practices, of which timing of harvest is very important (Shirtliffe et al., 2000; Walsh & Powles, 2014). Early harvesting always reduced seed shattering and increased harvestable seed yield, whereby the difference to the late harvest was highest for the F_2_-- population and lowest for the F_2_++ selected population. Hence, early timing of seed harvest is very suitable to enable high seed yields, regardless of the genetic material (Figure 4, Table 5). However, due to unfavorable weather conditions or other constraints, an early harvest is not always possible. Furthermore, early harvesting could have additional negative effects such as high moisture content in seeds resulting in higher costs for drying, or increased frequency of immature seeds resulting in lower germination rates (Hill & Watkin, 1975; Larson et al., 2020). Therefore, cultivars with reduced seed shattering are an important prerequisite for seed growers to ensure high seed yields also under non-optimal conditions.

Forage grass breeding mainly depends on recurrent phenotypic selection of populations (Posselt, 2010). Phenotypic selection is often based on selection among spaced plants and is especially useful for traits having a high heritability and good correlation between spaced plants and sward conditions. Earlier studies in perennial ryegrass showed a low correlation between seed yield in spaced plants and their offspring sown in plots (Bugge, 1987; Elgersma et al., 1994). Seed yield is influenced by seed shattering, but insufficient seed yield can have several other causes. For example, non-optimal pollination and fertilization, seed abortion or seed shattering are reported to be factors limiting seed yield (Falcinelli, 1999; Studer et al., 2008). Furthermore, seed yield responds differently depending on the competition in the growing environment (spaced plant with low space competition vs. plot trials with high space competition) (Elgersma, 1990b; Waldron et al., 2008). Especially for populations that already exhibit a low level of seed shattering, all these factors could have led to a low correlation between spaced plants and plot trials in these studies. Our study showed that the five populations divergently selected for seed shattering showed highly consistent results between spaced plant and plot trials (r = 1.00, *p* = 0.01). This indicates that selection for low seed shattering in spaced plants is transferable to sward conditions.

To conclude, our phenotyping method, which is based on bagging inflorescences after flowering followed by harvesting according to the sum of temperature, proved to be efficient to select for reduced seed shattering in spaced plants. To reduce the efforts needed for phenotyping, indirect improvement of seed shattering via selection for plants with higher mass of non-shattered seeds might be an alternative. Effects of selection were also significant in plot trials. Phenotypic selection in spaced plants is, therefore, effective to create cultivars with reduced seed shattering and increased harvestable yield under sward conditions, allowing farmers to ensure high seed yields also under non-optimal harvest conditions. The population selected for low seed shattering in this study is, therefore, a valuable genetic resource that can be directly used within breeding programs.

## CONFLICT OF INTEREST

The authors declare that the research was conducted in the absence of any commercial or financial relationships that could be construed as a potential conflict of interest.

## AUTHOR CONTRIBUTIONS

*Conceptualization*: Christoph Grieder and Roland Kölliker. *Methodology*: Christoph Grieder. *Formal analysis*: Jenny Kiesbauer. *Investigation*: Jenny Kiesbauer, Linda Schlatter, Maria Hug, Meril Sindelar and Jonathan Ohnmacht. *Data curation*: Jenny Kiesbauer, Jonathan Ohnmacht, Christoph Grieder. *Writing - original draft preparation*: Jenny Kiesbauer and Christoph Grieder. *Writing—review and editing*: Jenny Kiesbauer, Christoph Grieder, Roland Kölliker, Bruno Studer, Linda Schlatter. *Visualization*: Jenny Kiesbauer, Jonathan Ohnmacht. *Supervision*: Christoph Grieder, Roland Kölliker, Bruno Studer. *Funding acquisition*: Christoph Grieder. All authors have read and approved the final version of this manuscript.

## ACKNOWLEDGMENTS

The authors thank Peter Tanner, Simone Günter and Daniel Schmid from the Fodder Plant Breeding group at Agroscope for technical support in the field trials. This project received funding from Agroscope and Delley Seeds and Plants Ltd.

## DATA AVAILABILITY STATEMENT

The data presented in this study are openly available at zenodo.org (https://zenodo.org/record/10224788) (accessed on 30. November 2023)

## Notes

### Competing Interest Statement

The authors have declared no competing interest.

